# ADAM22 and ADAM23 modulate the targeting of the Kv1 channel-associated protein LGI1 to the axon initial segment

**DOI:** 10.1101/311365

**Authors:** Bruno Hivert, Laurène Marien, Komlan Nassirou Agbam, Catherine Faivre-Sarrailh

## Abstract

The distribution of voltage-gated potassium channels Kv1 at the axon initial segment (AIS), along the axon and at presynaptic terminals influences intrinsic excitability and transmitter release. Kv1.1/1.2 subunits are associated with cell adhesion molecules (CAMs), including Caspr2 and LGI1 that are implicated in autoimmune and genetic neurological diseases with seizures. In particular, mutations in the LGI1 gene cause autosomal dominant lateral temporal lobe epilepsy (ADTLE). In the present study, we used rat hippocampal neurons in culture to assess whether interplay between distinct Kv1-associated CAMs contributes to targeting at the AIS. Strikingly, LGI1 was highly restricted to the AIS surface when transfected alone, whereas the missense mutant LGI1^S473L^ associated with ADLTE was not. Next, we showed that ADAM22 and ADAM23 acted as chaperones to promote axonal vesicular transport of LGI1 reducing its density at the AIS. Moreover, live-cell imaging of fluorescently labelled CAMs indicated that LGI1 was co-transported in axonal vesicles with ADAM22 or ADAM23. Finally, we showed that ADAM22 and ADAM23 also associate with Caspr2 and TAG-1 to be selectively targeted within different axonal sub-regions. The combinatorial expression of Kv1-associated CAMs may be critical to tune intrinsic excitability in a physiological or an epileptogenic context.

## Introduction

Voltage-gated potassium Kv1 channels are concentrated at the axon initial segment (AIS) where they contribute to the control of neuronal excitability (Kole and Stuart, 2012; Rasband, 2010; Vacher and Trimmer, 2012). Kv1 channels co-purify with several cell adhesion molecules (CAMs) including Caspr2, TAG-1, LGI1 (Leucine-rich Glioma Inactivated 1) and ADAM (A Disintegrin And Metalloprotease) proteins, which may influence their positioning within the distinct axonal sub-regions. The importance of these CAMs in neuronal function is reflected by their implication in both genetic and autoimmune diseases associated with hyperexcitability and epilepsy (Kegel et al., 2012; Lai et al., 2010; Muona et al., 2016; Ohkawa et al., 2013; Rodenas-Cuadrado et al., 2014). Antibody-mediated limbic encephalitis which was first associated with voltage-gated potassium channels, has been mainly attributed to autoantibody binding to Caspr2 or LGI1 (Irani et al., 2010; Lancaster et al., 2011). Caspr2 is associated with TAG-1 at the juxtaparanodes of myelinated axons mediating axo-glial contacts and inducing the clustering of Kv1.1/1.2 to control the internodal resting potential (Poliak et al., 2003; Traka et al., 2003). In addition to juxtaparanodes, Caspr2 and TAG-1 are concentrated at the AIS of cortical and motor neurons, but they are not required for the recruitment of Kv1 at that site (Duflocq et al., 2011; Inda et al., 2006; Ogawa et al., 2008). In cultured hippocampal neurons, Kv1 channels are enriched at the AIS associated with TAG-1, whereas Caspr2 is targeted all along the axon (Pinatel et al., 2017). Other membrane proteins interacting with Kv1 channels may be localized at the AIS, including ADAM22 which is recruited at the AIS of cultured hippocampal neurons with PSD93 (Ogawa et al., 2010). However, while the voltage-gated sodium channels are known to be tethered by ankyrinG at the AIS (Rasband, 2010), the precise mechanisms implicated in the recruitment of the Kv1 complex at the AIS are still elusive.

LGI1 is a secreted glycoprotein consisting of leucine-rich and epitempin (EPTP) repeats that has been implicated in protein-protein interactions at the synapse, but was not described at the AIS until recently (Seagar et al., 2017). LGI1 interacts via its EPTP repeats with members of the ADAM family, including ADAM11, ADAM22 and ADAM23 (Fukata et al., 2006; Owuor et al., 2009; Sagane et al., 2008). LGI1 has been proposed to form a transynaptic complex with ADAM22 and ADAM23 controlling synaptic strength at excitatory synapses by regulating PSD-95 incorporation (Fukata et al., 2010; Lovero et al., 2015). LGI1 localized at the pre-synaptic terminals was reported to act as negative modulator of glutamate release, an effect which could be linked with pre-synaptic Kv1 (Boillot et al., 2016). In patients with autoimmune encephalitis, anti-LGI1 antibodies may disrupt interaction with ADAM proteins (Ohkawa et al., 2013). In addition, LGI1 is a monogenic human epilepsy-related gene, mutated in ADTLE (autosomal dominant temporal lobe epilepsy)(Gu et al., 2002; Kalachikov et al., 2002; Kegel et al., 2012; Morante-Redolat et al., 2002). LGI1 requires glycosylation to be secreted and most ADTLE mutations inhibit LGI1 secretion by preventing its proper folding. Interestingly, some mutations that do not inhibit secretion were found to impair interactions with ADAM22 and ADAM23 (Dazzo et al., 2016; Yokoi et al., 2015).

In the present study, we asked whether interplay between distinct sets of Kv1-associated CAMs affects axonal targeting in cultured hippocampal neurons. Our data indicate that these CAMs can be sorted together in axonally transported vesicles in cultured hippocampal neurons. We showed that ADAM22 and ADAM23 strongly increased the cell surface targeting of LGI1 and prevented its restricted expression in the AIS, in association with Kv1 channels. Importantly, we showed that the secreted missense mutants LGI1^S473L^ and LGI1^R474Q^ identified in ADTLE were not targeted to the AIS. This may induce perturbation of LGI1 function in tuning intrinsic excitability, thus contributing to epileptogenesis.

## Results

### ADAM22 and ADAM23 modulate the targeting of LGI1 at the AIS of hippocampal neurons

The Kv1 channels are known to associate with several membrane proteins, including ADAM22, ADAM23 and LGI1 at the presynaptic terminals. Here, we examined whether these CAMs may also act in interplay with the Kv1 complex at the AIS of hippocampal neurons in culture. LGI1 was recently reported to be enriched at the AIS of hippocampal CA3 neurons using immunofluorescence staining on brain sections (Seagar et al., 2017). In culture of hippocampal neurons, we found using the same anti-LGI1 mAb, that LGI1 was faintly expressed at the AIS surface at DIV8 (Fig. 1A). In contrast, LGI1 was present as small clusters mainly on the somato-dendritic compartment at DIV21, when the synaptic network is established (Fig. 1B). We analyzed the subcellular distribution of LGI1-GFP when transfected in hippocampal neurons either alone or co-transfected with its binding partners ADAM22 or ADAM23. We observed that LGI1-GFP was restricted at the AIS surface when transfected alone in hippocampal neurons at DIV8 as visualized using live immunostaining with anti-GFP antibody (Fig. 1C in green and 1H in red). The direct fluorescence of intracellular LGI1-GFP was faintly detected (Fig. 1H, green). Strikingly, co-transfection with ADAM22 or ADAM23 at DΓV8 strongly increased the cell surface expression of LGI1-GFP that was addressed to the somato-dendritic and axonal compartments as detected using live immunostaining with anti-GFP antibody (Fig. 1D, E, green; 1I, red). LGI1-GFP expressed alone displayed a fluorescence intensity AIS/axon ratio of 3.26 ± 0.23 (n=16). This ratio was significantly reduced to 1.38 ± 0.11 (n=24) and 1.15 ± 0.10 (n=19) when co-transfected with ADAM22 and ADAM23, respectively (Fig. 1J). As a control experiment we examined the polarized distribution of NrCAM-GFP since NrCAM does not belong to the Kv1 complex and is strongly recruited at the AIS by its AnkyrinG-binding motif (Davis et al., 1996). NrCAM-GFP displayed an AIS/axon ratio of 4.05 ± 0.6 (n=20) and 5.51 ± 0.9 (n=15) in the absence or presence of ADAM22, respectively (Fig. 1J). Therefore, the co-expression of ADAM22 did not modify the AIS distribution of NrCAM as illustrated in Fig. 1F, G.

**Figure 1:**
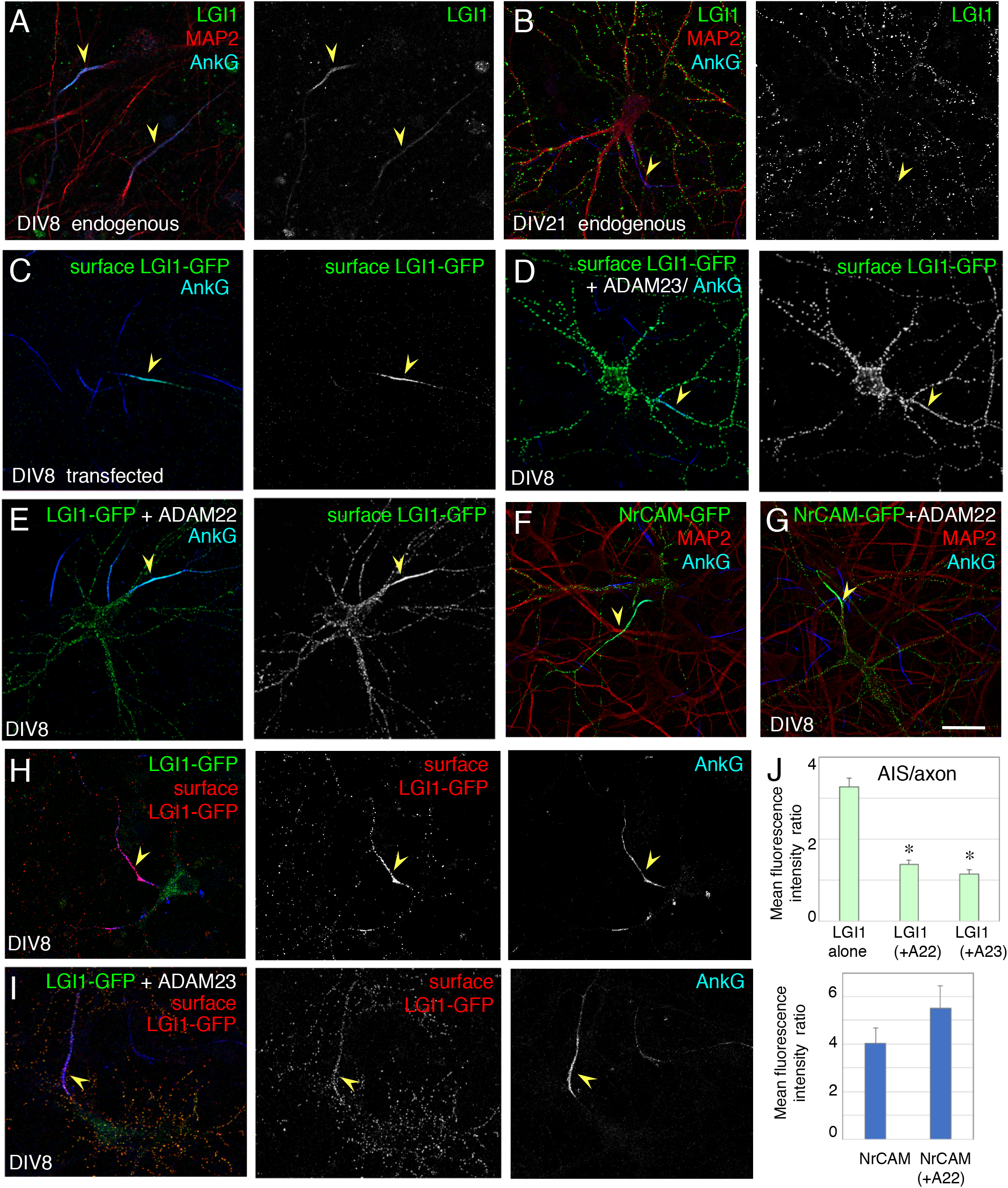
LGI1 enrichment at the AIS is modulated by its co-expression with ADAM22 and ADAM23 in cultured hippocampal neurons. (A, B) Hippocampal neurons at DIV8 or DIV21 were surface labeled using anti-LGI1 mAb (green), fixed permeabilized before immunostaining for AnkyrinG (blue) as a marker of the AIS (arrowheads) and MAP2 (red). Note that endogenous LGI1 is only detected at the AIS at DIV8 whereas it is mainly distributed as puncta on the somato-dendritic compartment at DIV21. (C-I) Hippocampal neurons were transfected at DIV8 with LGI1-GFP (C, H), LGI1-GFP and ADAM23 (D, I), LGI1-GFP and ADAM22 (E), NrCAM (F), or NrCAM and ADAM22 (G). (C-E) Neurons were surface labeled using anti-GFP antibodies (green), fixed and permeabilized before immunostaining for AnkyrinG (blue, arrowheads). Note that surface labeling of LGI1-GFP (C) is highly restricted to the AIS at DIV8. Co-transfection with ADAM22 (E) or ADAM23 (D) strongly enhances LGI1-GFP expression at the somatodendritic and axonal surface. Co-transfection with ADAM22 has no effect on NrCAM distribution at AIS (F, G). (H, I) Surface immunostaining of LGI1-GFP using AlexaFluor568 secondary antibodies (red) with direct imaging of LGI1-GFP fluorescence (green). Bar: 20 μm. (J) Ratios of fluorescence intensity between AIS and axon in neurons transfected at DIV8 with LGI1-GFP alone, NrCAM-GFP alone, or co-transfected with ADAM22 (A22) or ADAM23 (A23). *indicates significant differences by comparison with LGI1-GFP transfected alone for co-transfection with ADAM22 or ADAM23 (F(2, 56)= 55.68, ANOVA and p<0.01, Fisher test). The co-expression of ADAM22 did not modify the AIS distribution of NrCAM-GFP (Mann-Whitney test, p=0.13).

Next, the ADAM23 sequence was fused to mCherry at its C-terminal to facilitate its fluorescent detection (red) and it was surface labeled using rabbit anti-ADAM23 antibody (green) (Fig. S1B). ADAM23-mCherry exhibited a non-polarized surface expression when transfected alone (Fig. S1B) or co-transfected with LGI1-GFP (Fig. S1C). It displayed an AIS/axon ratio of 1.58 ± 0.13 (n=11) (Fig. S1D). Moreover, LGI1-GFP colocalized with ADAM23-mCherry in clusters both along dendrites (Fig. S1C’) and the axon labeled for AnkyrinG (Fig. S1C”).

### ADAM23 decreases the enrichment of LGI1-GFP co-localized with Kv1.2 channels at the AIS

We asked whether the recruitement of LGI1-GFP to the AIS may be correlated with the concentration of Kv1 channels at the AIS. In contrast to the early appearance of AnkyrinG, NrCAM and voltage-gated sodium channels, the Kv1.1/1.2 channels are tethered at the AIS of cultured hippocampal neurons only after 10 days in vitro (Sanchez-Ponce et al., 2012; Vacher and Trimmer, 2012; Vacher et al., 2011). We analyzed neurons transfected with LGI1-GFP or co-transfected with LGI1-GFP and ADAM23-mCherry at DIV14 and the expression of endogenous Kv1.2 was measured at the AIS and along the axon (Fig. 2). When LGI1-GFP was transfected alone, it was found enriched at the AIS and co-localized with Kv1.2 (Fig. 2A). The ratio of the mean fluorescence intensity at the AIS versus the axon was 2.82 ± 0.3 for LGI1-GFP and 2.47 ± 0.24 for Kv1.2. Individual values (n=13) were plotted showing that the AIS/axon ratio for LGI-GFP may be correlated with the one of the Kv1.2 channels (Fig. 2C). In contrast, when expressed with ADAM23-mCherry, LGI1-GFP was not anymore enriched at the AIS of co-transfected neurons (Fig. 2B, D). This result suggests that ADAM23 removes LGI-GFP from the AIS of the co-transfected neuron, which is not anymore associated in a complex with the Kv1 channels.

**Figure 2:**
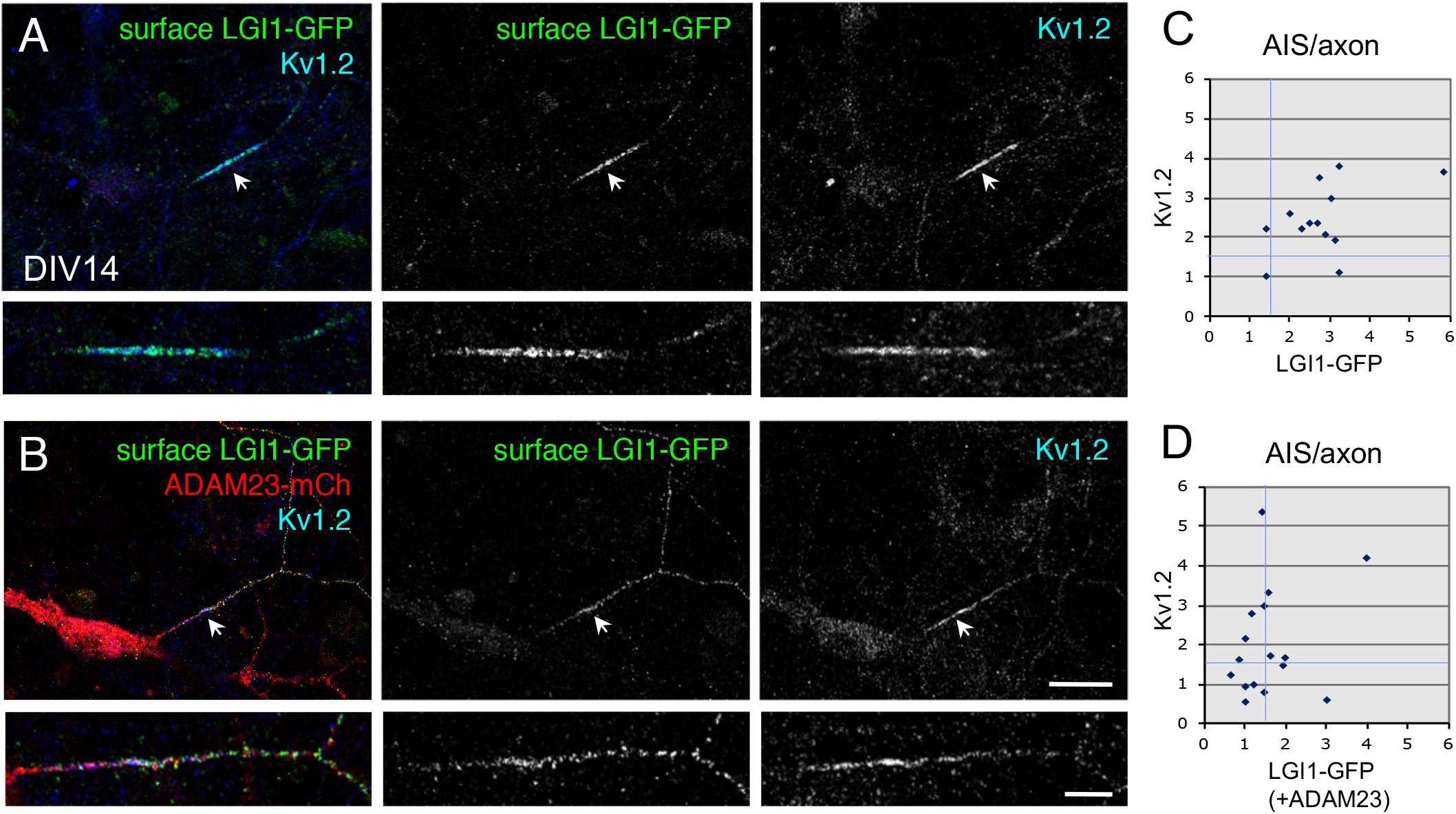
The AIS enrichment in LGI1-GFP correlated with AIS expression of endogenous Kv1.2. DIV13 hippocampal neurons were transfected with LGI1-GFP (A) or with LGI1-GFP and ADAM23-mCherry (B). Neurons were surface labeled for GFP (green), fixed, permeabilized and immunostained for Kv1.2 (blue). Note that LGI1-GFP is colocalized with Kv1.2 at the AIS in a neuron transfected with LGI1-GFP alone, but not in a neuron co-transfected with LGI1-GFP and AD AM23-mCherry. The AIS is indicated with white arrows. (C, D) The AIS/axon ratios of fluorescence intensity for LGI1-GFP and Kv1.2 were plotted for individual neurons when transfected with LGI1-GFP (C) or co-transfected with LGI1-GFP and ADAM23-mCherry (D). Bar: 20 μm, in inset: 5 μm.

### ADAM22 and ADAM23 promote ER export of LGI1

Our data indicated that the ADAM proteins may favor surface expression of LGI1 either by stabilizing the secreted glycoprotein at the cell membrane or by promoting its export along the secretory pathway. We analyzed whether the ADAM proteins may be implicated in the intracellular trafficking of LGI1 using transfected HEK cells. We observed that LGI1-GFP was strongly retained in the ER (green) when transfected alone in HEK cells and was poorly detected at the cell surface using live immunolabeling for GFP and Alexa647 secondary antibody (blue) (Fig. 3A). In contrast, LGI1-GFP co-expressed with ADAM23-mCherry was faintly detected in the ER and strongly labeled as clusters at the cell surface (blue) (Fig. 3C). When co-transfected with ADAM22-mCherry, LGI1-GFP was also detected at the cell surface (Fig. 3B). Thus, we deduced that the association with ADAM proteins could favor the transport permissive conformation of LGI1.

**Figure 3:**
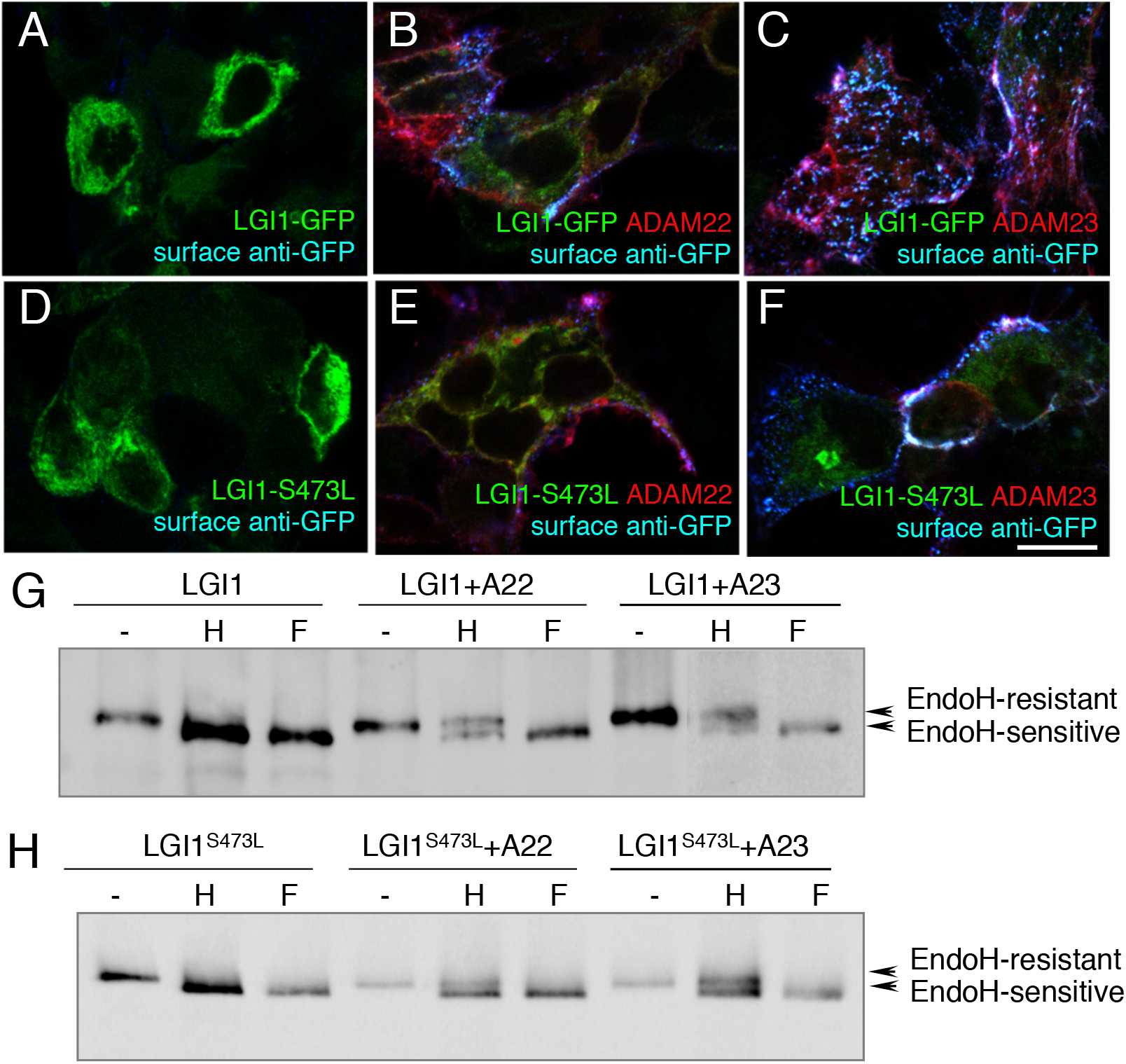
ADAM22 and ADAM23 promote ER exit and N-glycan maturation of LGI1 and LGI1^S473L^. HEK cells (A-F) were transfected with LGI1-GFP alone (A), or co-transfected with ADAM22-(B) or ADAM23-mCherry (C), transfected with LGI1^S473L^ alone (D), or cotransfected with ADAM22-(E) or ADAM23-mCherry (F). The fluorescence for GFP was directly imaged (green) to visualize the intracellular pool of LGI1 whereas the surface pool was labeled using anti-GFP antibody (blue). Note that ADAM22 and ADAM23 promote ER exit and surface expression of LGI1 and LGI1^S473L^. (G, H) HEK cells were transfected with LGI1-GFP (G) or LGI1^S473L^ (H) alone or co-transfected with ADAM22-mCherry or ADAM23-mCherry. After cell lysis, wild type or mutated LGI1 was immunoprecipitated using anti-GFP mAb and incubated at 37°C for 3 h without (-) or with Endo H (H) or PNGase F (F). Western blotting with anti-GFP mAb shows that LGI1-GFP migrates as a doublet when co-transfected with ADAM22 or ADAM23 and incubated with Endo H. Note that the lower band corresponding to the Endo H-sensitive gly coform migrates as LGI1-GFP treated with PNGase F. The higher band, which is Endo H-resistant migrated as untreated LGI1-GFP. The LGI1^S473L^ mutant also displayed Endo H-sensitive and resistant glycoforms. Bar in A-F: 10 μm.

Next, we examined the N-glycosylation processing of LGI1-GFP when expressed alone or in association with ADAM22 or ADAM23 in HEK cells. We analyzed the glycosylation pattern of LGI1-GFP using endoglycanase H (Endo H), which digests only immature ER-type N-glycans bearing high-mannose. N-glycosidase F (PNGase F) was used to remove all the N-glycans. LGI1-GFP contained Endo H-sensitive carbohydrates when transfected alone (Fig. 3G). In contrast, when co-transfected with ADAM22 or ADAM23, two bands of LGI1-GFP were detected after treatment with Endo H, the higher band being Endo H-resistant and the lower band Endo H-sensitive (Fig. 3G). This result indicates that ADAM22 and ADAM23 favor ER exit of LGI1 likely by acting as chaperone-like proteins through the ER quality - control system in HEK cells. In accordance with the results of Yokoi et al. (2015), we observed that Endo H digestion produced a mobility shift of LGI1-GFP while PNGase F treatment induced a further mobility shift of the same pool, as an indication that LGI1 can be partially processed in the absence of ADAM proteins.

### The secreted mutants LGI1^S473L^ and LGI1^R474Q^ are not addressed at the AIS

Human epilepsy-related missense mutations of LGI1 have been reported and classified as secretion-defective or secretion-competent mutations. Among this last category, LGI1^S473L^ which is mutated in the EPTP domain, displays reduced binding ability for ADAM22 but not ADAM23 (Yokoi et al., 2015). Therefore, we asked whether LGI1^S473L^ could be properly addressed to the AIS of hippocampal neurons when expressed alone or in combination with ADAM22 or ADAM23. First, we analyzed the processing of LGI1^S473L^ in HEK cells and observed that both ADAM22 and ADAM23 induced its cell surface expression (Fig. 3D-F). Western blotting experiments indicated that LGI1^S473L^ displayed Endo H-sensitive N-glycans when expressed alone in HEK cells, showing the same mobility shift as the wild type LGI1-GFP after Endo H or PNGase F treatment (Fig. 3H). LGI1^S473L^ co-expressed with ADAM22 or ADAM23 displayed both Endo H-resistant and Endo H-sensitive glycoforms (Fig. 3H). Thus, LGI1^S473L^ still had the capacity of interacting with ADAM proteins, which enhanced its ER exit and processing with complex N-glycans.

Strikingly, when transfected in hippocampal neurons at DIV8, LGI1^S473L^ was not enriched at the AIS (Fig. 4C) in contrast to wild type LGI1-GFP (Fig. 4A). The mean AIS/axon ratios were 3.62 ± 0.39 (n=20) for LGI1-GFP and 1.18 ± 0.09 (n=17) for LGI1^S473L^. Co-transfection with ADAM22- or ADAM23-mCherry strongly enhanced the surface labeling of LGI1^S473L^ that became unpolarized at the neuronal cell surface (Fig. 4G, H). These data indicate that the binding activity of LGI1^S473L^ for ADAM22 and ADAM23 allows its transport-permissive conformation in hippocampal neurons. Next, we tested the distribution of two other secreted mutants of LGI1, LGI1^R407C^ and LGI1^R474Q^ when transfected in DIV8 hippocampal neurons. As expected, the R474Q mutation, which is adjacent to the S473L mutation also prevented the AIS enrichment of LGI1 (Fig. 4D, J). In contrast, the LGI1^R407C^ was detected enriched at the AIS of hippocampal neurons as observed for the wild type LGI1 (Fig. 4B, J). The mean AIS/axon ratios were 1.55 ± 0.13 (n=17) for LGI1^R474Q^ and 2.72 ± 0.19 (n=19) for LGI1^R407C^. Co-transfection with ADAM22- or ADAM23-mCherry strongly enhanced the neuronal cell surface expression of LGI1^R474Q^ (not shown) and LGI1^R407C^ (Fig. 4E, F). In conclusion, two missense mutations located in the EPTP6 domain, the S473L and R474Q mutations both impair the recruitment of LGI1 at the AIS as a possible pathogenic mechanism. In contrast, the mutation R407C in the EPTP4 domain has no effect on AIS targeting (Fig. 4I). The association of the three LGI1 mutants with co-transfected ADAM22 and ADAM23 is sufficient to increase their expression at the neuronal cell surface.

**Figure 4:**
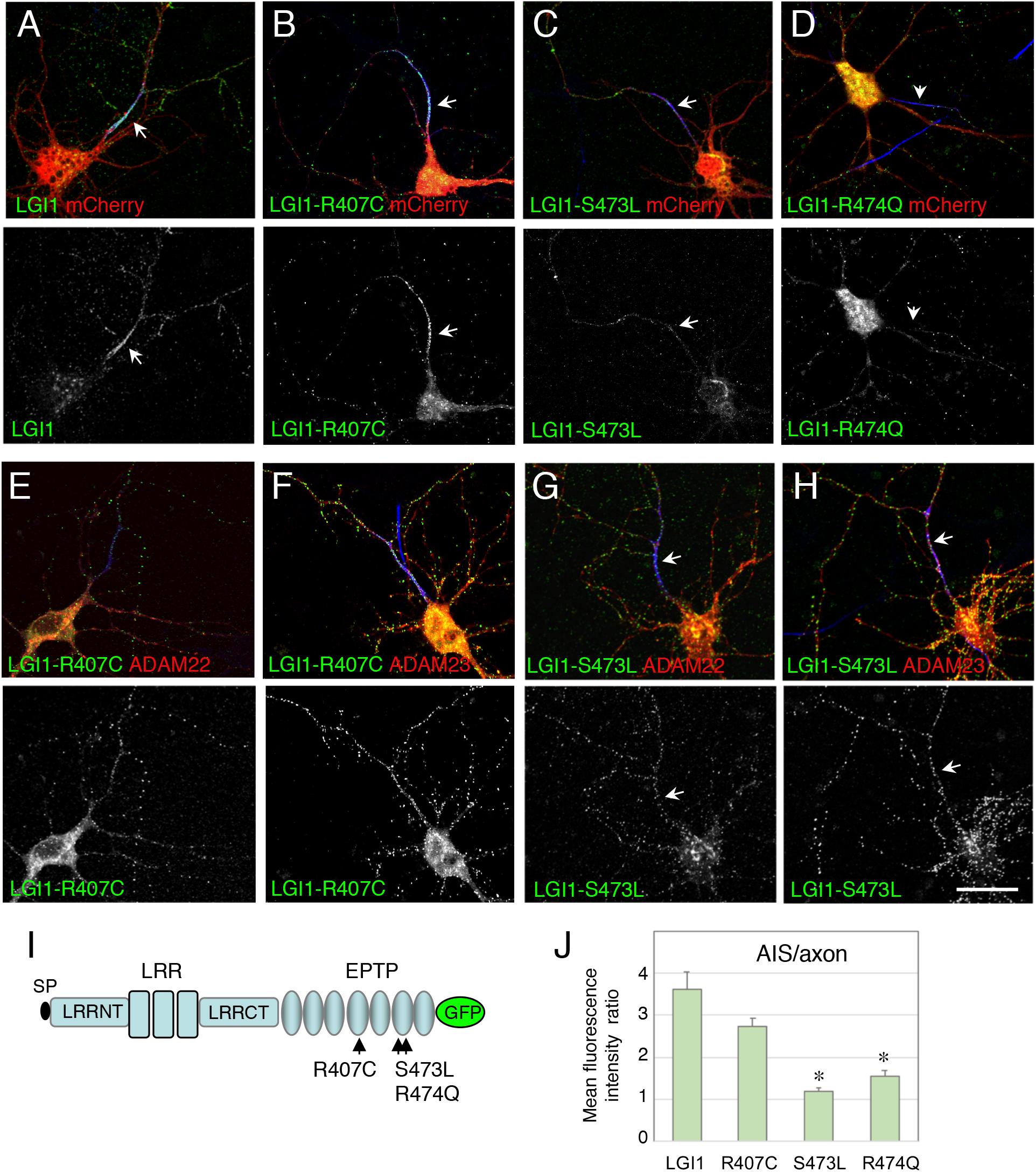
The variants of LGI1 associated with epilepsy are differentially recruited at the AIS of hippocampal neurons. DIV8 hippocampal neurons were transfected with LGI1-GFP (A), LGI1^R407C^ (B), LGI1^S473L^ (C), LGI1^R474Q^ (D). Note that both LGI1^S473L^ and LGI1^R474Q^ are not enriched at the AIS in contrast to wild type LGI1 and LGI1^R407C^. LGI1^R407C^ or LGI1^S473L^ were co-transfected with ADAM22-(E, G) or ADAM23-mCherry (F, H). When co-transfected with ADAM22- or ADAM23-mCherry, the neuronal surface expression of all the variants is strongly increased. (I) Schematic representation of LGI1 with the point mutations localized in the EPTP domains. Leucine Rich Repeats (LRR), N-terminal (NT) and C-terminal (CT), and EPTP domains. (J) Ratios of fluorescence intensity between AIS and axon in neurons transfected at DIV8 with LGI1-GFP, LGI1^R407C^, LGI1^S473L^ or LGI1^R474Q^. indicates significant differences by comparison with wild type LGI1-GFP (P<0.0001, Mann-Whitney test). Bar: 20 μm.

### Axonal transport of ADAM22, ADAM23 and LGI1

We next investigated whether LGI1 could be associated with ADAM proteins in axonal transport vesicles. First, we performed time-lapse imaging of neurons transfected at DIV8 with ADAM22-mCherry or ADAM23-mCherry to get insights into their axonal targeting mechanisms. The axon was clearly identified on the basis of its length (and was strongly enriched in transport vesicles by comparison with dendrites). In addition, live immunolabeling of Neurofascin-186 (blue) was used to precisely localize the AIS after time-lapse recording (Fig. 5A, B). In neurons that were transfected with ADAM22-mCherry, we observed that labeled vesicles were axonally transported in the anterograde and retrograde directions with a maximal velocity (Vm) of 0.69 ± 0.1 and 0.51 ± 0.06 μm/s, respectively (Table S1, Movie 1). In ADAM23-mCherry-transfected neurons, labeled vesicles were transported in the anterograde and retrograde directions with a Vm of 0.99 ± 0.11 and 0.53 ± 0.06 μm/s, respectively (Table S1, Movie 2). However, ADAM22 vesicles moved bidirectionally in most neurons (Table S1, n=10 neurons) whereas by contrast ADAM23 vesicles were mostly transported in the anterograde direction (65.2 ± 9.5 % of displacements, n=9 neurons) as indicated by kymograph analysis (Fig. 5A’, B’).

**Figure 5:**
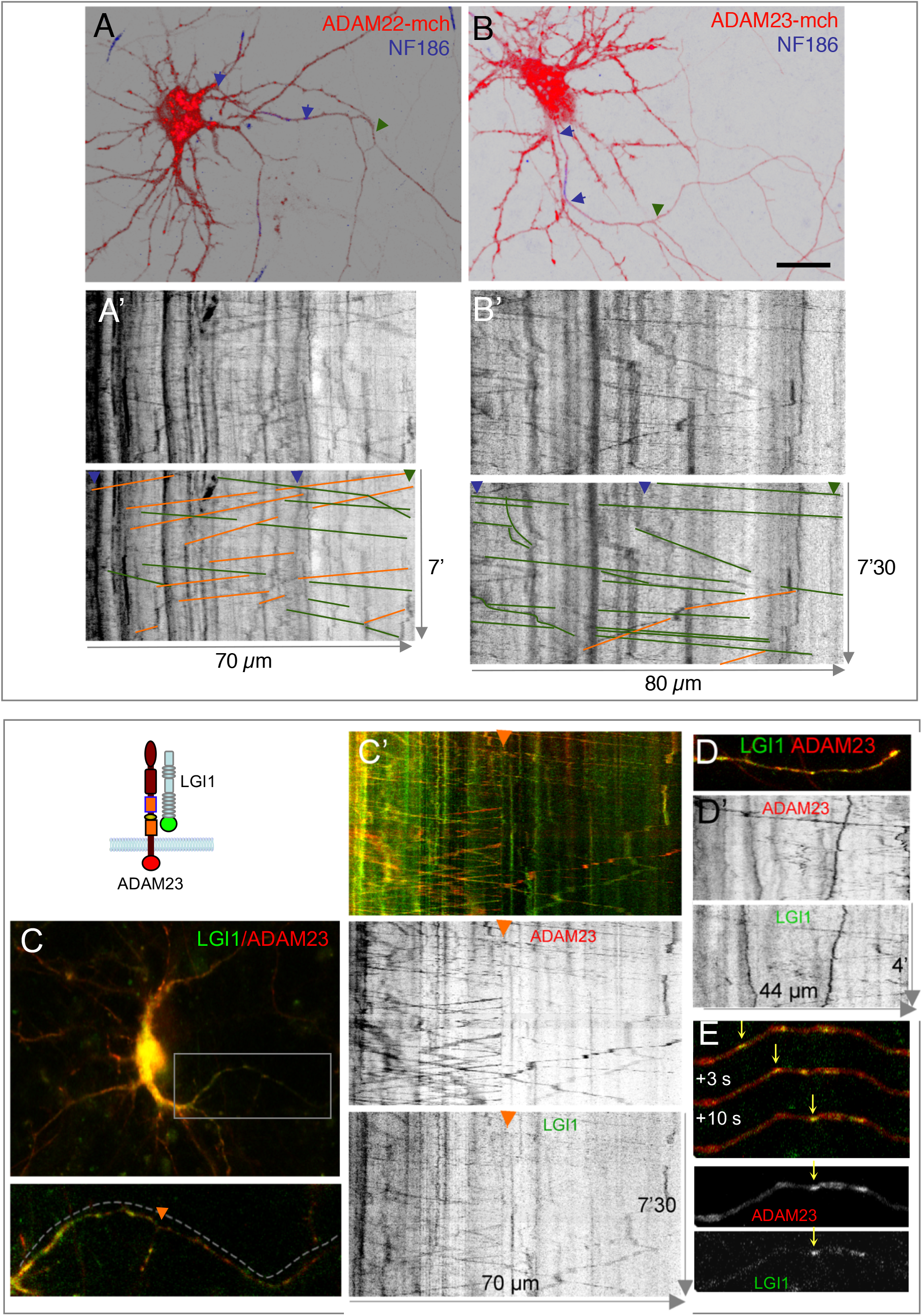
LGI1 and ADAM23 are co-localized in transport vesicles. (A, B) Hippocampal neurons were transfected at DIV8 with ADAM22-mCherry (A), or ADAM23-mCherry (B). Streamed time-lapse images of axonal transport vesicles were acquired at 1 frame per 1.5 s. Live immunostaining with Alexa647-coupled anti-Neurofascin186 was performed to determine AIS location (blue) limited with blue arrowheads. (A’, B’) Corresponding kymographs with axonal length in x-axis and time in y-axis. Anterograde and retrograde events are underlined with green and orange traces, respectively. The maximal velocity was measured for each transport sequence. Note that transport events were mostly in the retrograde direction for ADAM22 whereas they were mostly in the anterograde direction for ADAM23. See the corresponding Movies 1 and 2. (C-E) Hippocampal neurons were co-transfected with LGI1-GFP and ADAM23-mCherry at DIV8. Live-cell recording of proximal (C) and distal (D, E) axons. The orange arrowhead labels a proximal axon bifurcation in C, C’. (C’, D’) The corresponding kymographs show overlapping trajectories of vesicles labelled for LGI1-GFP and ADAM23-mCherry. Comparison of traces for transport events along the axon indicated that a number of vesicles were colabeled with LGI1-GFP and ADAM23-mCherry moving both in anterograde and retrograde directions. (E) Time-lapse sequence showing a moving vesicle that contains both LGI1-GFP and ADAM23-mCherry indicated with arrows. Movies 3 and 4 show time-lapse recordings of proximal and distal axons, respectively. Bar: A-C, 20 μm.

Next, we performed two-color time-lapse imaging of neurons co-transfected with ADAM23-mCherry and LGI1-GFP to visualize their axonal transport (Fig. 5C-E). We found that most of the axonal transport vesicles were co-labeled for LGI1 and ADAM23. Kymograph analysis of transport events indicated that double-labeled vesicles moved bidirectionally as illustrated in proximal (Fig. 5C, C’) or distal (Fig. 5D-E) axonal regions (Movies 3 and 4). Vm for the anterograde and retrograde transports were 0.87 ± 0.09 and 0.86 ± 0.08 μm/s, respectively. We observed mostly anterograde events (74 ± 4.7% of displacements, n=4 neurons), as occurring in neurons transfected with ADAM23 alone. In neurons co-transfected with ADAM22-mCherry and LGI1-GFP, we also observed axonal vesicles co-labeled for both CAMs and mostly retrogradely transported (Fig. S2, n=3 neurons). Vesicles labelled for LGI1-GFP were not easily detected when transfected alone, further indicating that LGI1 may require co-expression with ADAM22 or ADAM23 for its proper trafficking and axonal transport.

### Biochemical interactions of ADAM22 and ADAM23 with the Kv1-associated CAMs, TAG-1 and Caspr2

Apart from LGI1 and ADAM proteins, another set of CAMs including TAG-1 and Caspr2, has been reported to associate with Kv1 channels at discrete regions of the axon and we investigated whether these different CAMs may interplay for targeting. The biochemical interactions between LGI1 and ADAM22 or ADAM23 have been well documented: LGI1 interacts via its EPTP repeats with several members of the ADAM family, including ADAM22, and ADAM23 as reported using cell binding assays and co-immunoprecipitation experiments (Fukata et al., 2006; Owuor et al., 2009; Sagane et al., 2008). We performed co-imunoprecipitation experiments from transfected HEK cells to investigate whether ADAM22 and ADAM23 may also interact with the other CAMs of the Kv1 complex, TAG-1 and Caspr2. As shown in Fig. 6A, using co-immunoprecipitation with anti-mCherry or anti-GFP antibodies, LGI1-GFP but not TAG-1-GFP could associate in complex with ADAM23-mCherry. In addition, we showed that HA-tagged Caspr2 was co-immunoprecipitated with ADAM23-mCherry using anti-mCherry antibody (Fig. 6B). Moreover, Caspr2 deleted from its cytoplasmic tail (Caspr2Δcyt) was also efficiently co-immunoprecipitated with ADAM23, demonstrating that these membrane proteins interact via their ectodomains (Fig. 6B). Conversely, ADAM23 was not precipitated with Caspr2 when using anti-HA antibody. It may be noted that mCherry is fused at the C-terminus of ADAM23 cytoplasmic tail whereas HA is placed at the N-terminal region of Caspr2, so that only the anti-HA antibody may possibly interfere with the binding between CAM ectodomains.

**Figure 6:**
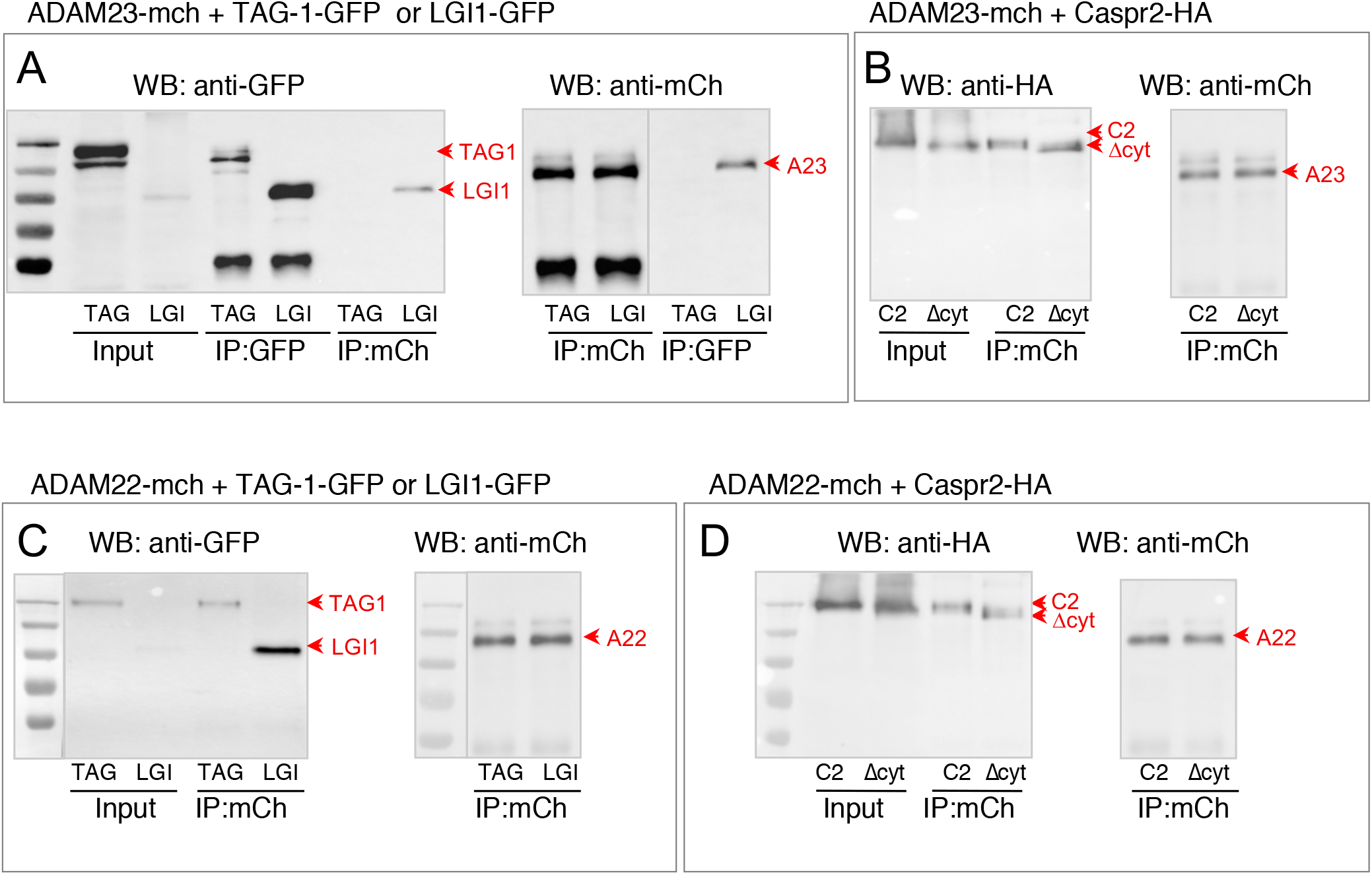
Biochemical analysis of the interaction between ADAM proteins and LGI1, TAG-1 or Caspr2. Co-immunoprecipitation experiments from HEK cells transfected with ADAM23-mCherry (A, B) or ADAM22-mCherry (C, D) and TAG-1-GFP or LGI1-GFP (A, C), or Caspr2-HA constructs either full-length (C2) or deleted from the cytoplasmic tail (Δcyt) (B, D). Immunoprecipitation of ADAM22- or ADAM23-mCherry was performed using rabbit anti-mCherry (IP mCh). Immunoprecipitation of TAG-1-GFP or LGI1-GFP was performed with mouse anti-GFP mAb (IP GFP). After Western blotting, TAG-1 and LGI1 constructs were detected using mouse anti-GFP mAb, ADAM proteins using rabbit anti-mCherry, and Caspr2-HA constructs using rat anti-HA mAb in the lysates (input) and immunoprecipitates. LGI1-GFP was co-immunoprecipitated with ADAM22 and ADAM23 (A, C). TAG-1 was co-immunoprecipitated with ADAM22 (C) but not with ADAM23 (A). Both full-length Caspr2 and Caspr2 deleted from its cytoplasmic tail were co-immunoprecipitated with ADAM22 and ADAM23 (B, D). Experiments were performed in triplicate.

Next, we showed that LGI1-GFP and also TAG-1-GFP were co-immunoprecipitated with ADAM22-mCherry (Fig. 6C). Caspr2 and Caspr2Δcyt were similarly co-immunoprecipitated with ADAM22-mCherry using anti-mCherry antibody (Fig. 6D). Thus, when expressed in HEK cells, ADAM22 and ADAM23 display the capability to selectively associate with multiple CAMs related to the Kv1 complex.

### The axonal targeting of ADAM23 is modulated by its co-expression with TAG-1 and Caspr2

We asked whether the ADAM proteins could interfere with the axonal targeting of the two components of the Kv1 complex, Caspr2 and TAG-1. We recently showed that both endogenous and transfected TAG-1 and Caspr2 are differentially distributed in cultured hippocampal neurons: TAG-1 is enriched at the AIS whereas Caspr2 is evenly localized along the axon (Pinatel et al., 2017). Neurons were co-transfected at DIV13 with TAG-1-GFP and ADAM23 and surface labeled using anti-GFP and anti-ADAM23 antibodies. Strikingly, we observed that ADAM23 colocalized with TAG-1 at the neuronal surface and was enriched at the AIS (Fig. S3A). In contrast, ADAM23 was faintly detected along the axonal surface when co-transfected with Caspr2-HA (Fig. S3B). Since Caspr2 is strongly internalized in the somato-dendritic compartment (Bel et al., 2009), we analyzed whether ADAM23 could be associated with Caspr2 in endocytic vesicles. Indeed ADAM23 was co-localized with Caspr2 in intracellular vesicles using immunostaining on fixed and permeabilized neurons (Fig. S3C). Next, neurons were co-transfected with TAG-1-GFP or Caspr2-HA, and ADAM23-mCherry to better visualize ADAM23 and perform quantitative analysis. The direct fluorescence of ADAM23-mCherry was strongly detected at the AIS in DIV14 hippocampal neurons, when co-expressed with TAG-1-GFP (Fig. 7A), but not with Caspr2-HA (Fig. 7B). The AIS/axon ratio of ADAM23-mCherry expressed alone or co-transfected with Caspr2 was 1.5 ± 0.1 (n=9) and 1.3 ± 0.1 (n=16), respectively. ADAM23 was significantly enriched at the AIS when co-transfected with TAG-1 with an AIS/axon ratio of 2.0 ± 0.2 (n=11) (Fig. 7C).

**Figure 7:**
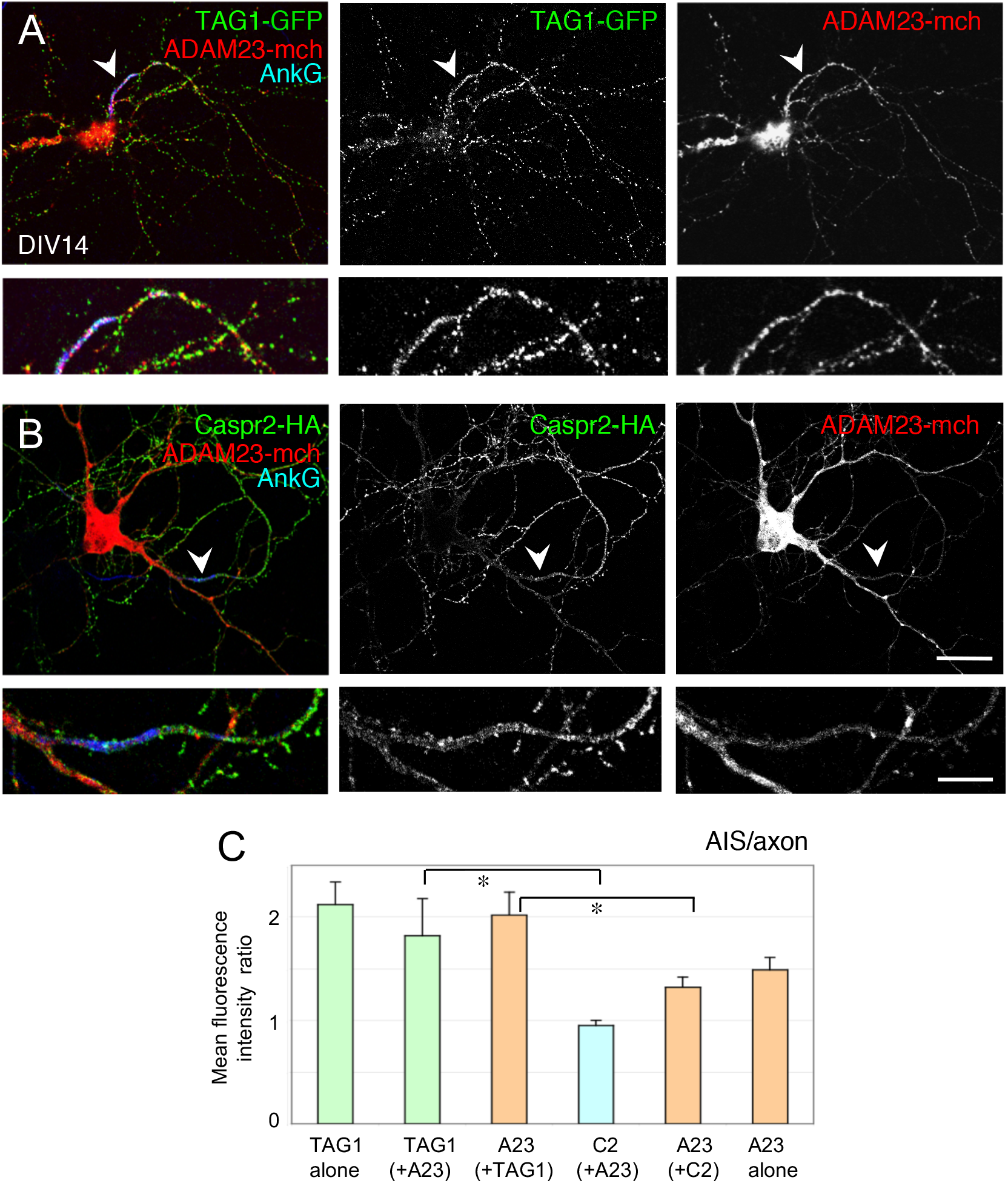
ADAM23-mCherry is co-targeted with TAG-1-GFP at the AIS of hippocampal neurons. Hippocampal neurons were co-transfected at DIV13 with ADAM23-mCherry and TAG-1-GFP (A), or with Caspr2-HA and ADAM23-mcC erry (B, C). Neurons were surface labeled with mouse anti-GFP mAb (A, green) or rat antiHA (B, green) antibodies and the fluorescence of ADAM23-Cherry was directly imaged (red). Cells were fixed and permeabilized before immunostaining for AnkyrinG (blue, arrowheads). Note that ADAM23-mCherry was enriched and colocalized with TAG-1 at the AIS (A, inset) whereas it was faintly detected at the AIS when co-transfected with Caspr2-HA (B). (C) Ratios of fluorescence intensity between AIS and axon for TAG-1-GFP, ADAM23-mCherry (A23), and Caspr2 (C2), either transfected alone or co-transfected. *indicated significant differences using the non-parametric Mann-Whitney test (P<0.01).

Using time-lapse live-cell imaging, we analyzed whether TAG-1, Caspr2 and ADAM23 may be sorted within the same axonal transport vesicles (Fig. 8). We observed vesicles co-labeled with ADAM23-mCherry and TAG-1-GFP moving along the axon (Fig. 8A-E; Movies 5 and 6). Kymograph analysis of transport events indicated that these vesicles moved in the anterograde and retrograde directions with a Vm of 1.1± 0.3 μm/s and 0.7± 0.1 μm/s, respectively (n=8 neurons) (Table S2). We determined that vesicles co-labeled for TAG-1 and ADAM23 moved mostly retrogradely as previously observed for TAG-1(Pinatel et al., 2017). Therefore, ADAM23-mCherry was preferentially transported anterogradely when expressed alone (65 ± 9 % of anterograde displacements, Table S1, Fig. 5B, B’). In contrast, it moved mostly retrogradely when associated with TAG-1 (only 34 ± 6 % of anterograde displacements, Table S2, Fig. 8B, D). The retrograde vesicles labelled for TAG-1 and ADAM23 are likely endosomes and might provide AIS enrichment at the steady-state by recycling the distal axonal membrane.

**Figure 8:**
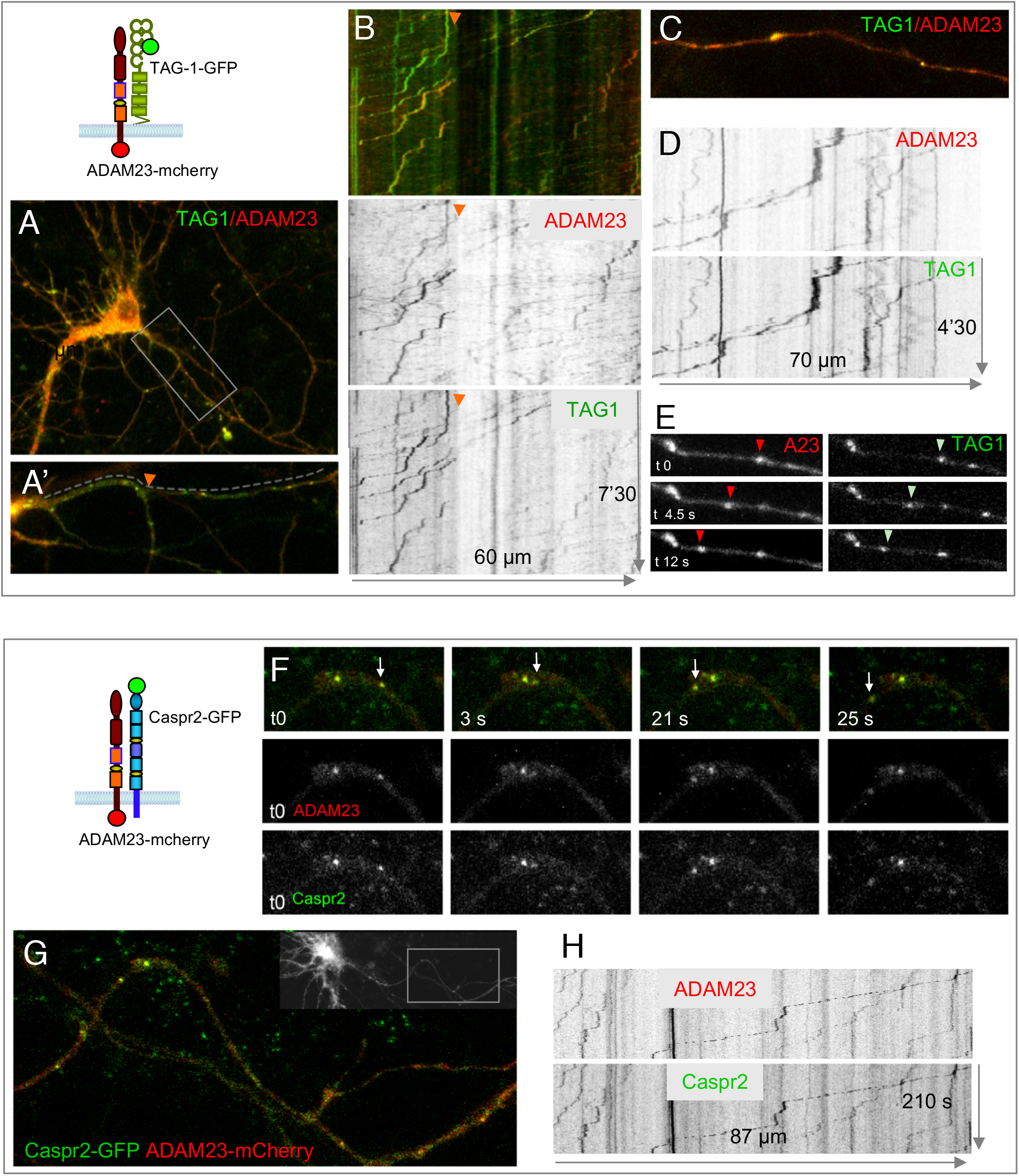
ADAM23-mCherry can be sorted with TAG-1-GFP or Caspr2-GFP within the same axonal transport vesicles. (A-E) Neurons co-transfected at DIV8 with TAG-1-GFP and ADAM23-mCherry. Stream time-lapse images of transport vesicles were acquired at 1 frame per 1.5 s in proximal (A, A’) and distal (C) axonal regions. (B, D) The corresponding kymographs showed overlapping trajectories of vesicles labeled for TAG-1-GFP and ADAM23-mCherry. The horizontal arrow indicates the distance along neurite and orientation. The orange arrowhead indicates the position of the axonal bifurcation (A’, B). (E) Example of a sequence showing a moving colabeled vesicle. The vesicles moved mostly in the retrograde direction. Movies 5 and 6 show time-lapse recording of these axons. (G-H) Hippocampal neuron transfected at DIV8 with Caspr2-GFP and ADAM23-mCherry. Stream time-lapse images of axonal transport vesicles were acquired at 1 frame per 1.5 s for 210 s in the region shown in the inset in G. (F) Time-lapse sequence showing a moving vesicle that contains both Caspr2-GFP and ADAM23-mCherry. (H) Corresponding kymographs showing overlapping trajectories of vesicles labeled for Caspr2-GFP and ADAM23-mCherry. Movie 7 shows time-lapse recording of this axon.

Next, we examined axonal transport in neurons co-expressing ADAM23-mCherry and Caspr2-GFP (Fig. 8F-H, Movie 7). Kymograph analysis of transport events indicated that these vesicles moved in the anterograde and retrograde directions with a velocity of 1.4 and 1 μm/s, respectively (Fig. 8H).

In conclusion, we showed that the combinatorial expression of CAMs associated with the Kv1 complex in hippocampal neurons may impact their respective subcellular distribution along the axon. ADAM23 was differentially distributed when co-expressed with TAG-1 or with Caspr2 being either enriched at the AIS or homogenously expressed all along the axon, respectively. Moreover our data indicate that these CAMs can be sorted together in axonally transported vesicles in cultured hippocampal neurons.

## Discussion

In the present study, we analyzed the axonal targeting of CAMs associated with Kv1 channels in cultured hippocampal neurons. We showed that LGI1 is enriched at the AIS in cultured hippocampal neurons and that the human mutations LGI1^S473L^ and LGI1^R474Q^ associated with epilepsy impair its trapping at the AIS as a possible pathogenic mechanism. The co expression of ADAM22 or ADAM23 strongly modulates the targeting of LGI1, which is decreased at the AIS and increased at the overall neuronal cell surface. We demonstrated that association with ADAM proteins is not only required for the anchoring of LGI1 at the cell surface, but is also implicated in its trafficking including ER exit, N-glycosylation processing, and axonal vesicular transport. In addition, an interplay was evidenced between ADAM proteins and other CAMs of the Kv1 complex, TAG-1 and Caspr2, that may be implicated in their selective targeting at the AIS.

We observed that LGI1 is tethered at the AIS when transfected in cultured hippocampal neurons. Next, we found that its co-expression with ADAM22 and ADAM23 strongly modulates the subcellular distribution of LGI1. Co-transfection with the ADAM proteins strongly increases the surface expression of LGI1 at the somato-dendritic and axonal compartments. As a consequence, the enrichment of LGI1 at the AIS becomes reduced in neurons overexpressing ADAM22 and ADAM23. Thus, the association of LGI1 with the Kv1 channels at the AIS, along the axon or at the synaptic terminals, may be modulated depending on its association with ADAM22 or ADAM23. ADAM22 has been described at the AIS of cultured hippocampal neurons colocalized with Kv1 channels and PSD93, but it is not required for the AIS clustering of Kv1.2 as indicated by the phenotype of *adam22^-/-^* mice (Ogawa et al., 2010). However reciprocally, high expression of ADAM22 or ADAM23 may remove LGI1 associated in a complex with the Kv1 channels from the AIS towards the axonal terminals, through a competition with ADAM22 tethered at the AIS.

LGI1 has been reported to act presynaptically as a negative regulator of excitatory transmission in early postnatal stages, possibly through Kv1-mediated modulation of synaptic release (Boillot et al., 2016). More recently, LGI1 was shown to localize at the AIS of CA3 hippocampal neurons regulating action potential firing by controlling the density of Kv1 channels (Seagar et al., 2017). LGI1 knock-out mice display a downregulation of the expression of Kv1.1 and Kv1.2 via a post-translational mechanism that may contribute to epileptogenesis. Interestingly, we show that the human mutations LGI1^S473L^ and LGI1^R474Q^ associated with epilepsy impairs LGI1 trapping at the AIS of hippocampal neurons as a possible pathogenic mechanism. In contrast, the LGI1^R407C^ mutant is properly recruited at the AIS indicating that another function of LGI1 is affected in this secreted mutant. Interestingly, the LGI1^S473L^ mutant is properly recruited into the synaptosomal fractions from transgenic mice (Yokoi et al., 2015), suggesting multiple roles of LGI1 according to its localization.

The ADAM proteins may act either as chaperones promoting LGI1 export or as anchors to stabilize the secreted LGI1 protein at the neuronal cell surface, which are indeed two non exclusive processes. We demonstrated that LGI1 cell surface targeting is strongly increased by its binding to ADAM22 or ADAM23 both in HEK cells and in neurons. Second, a distinct carbohydrate processing is observed when LGI1 is co-expressed with ADAM22 or ADAM23 in HEK cells with a doublet of Endo H-sensitive and Endo H-resistant glycoforms while an intermediate shift after Endo H treatment is detected when LGI1 is expressed alone. These results indicate that binding with ADAM proteins may induce a conformational switch further enhancing the ER exit of LGI1. LGI1 may bear distinct carbohydrates when associated or not with ADAMs, and when targeted at the synapses or at the AIS.

In addition, most of the missense mutations identified in ADTLE were classified as secretion-defective mutations, indicating that this genetic disorder can be a conformational disease (Yokoi et al., 2015). The LGI1^S473L^ mutant that is secreted, was shown to exhibit defective binding activity for ADAM22 using tandem affinity purification from transgenic mouse brain (Yokoi et al., 2015). A consequence of this reduced binding activity might be to interfere with the optimal transport-permissive conformation of the mutant protein. However, we show here that LGI1^S473L^ sufficiently associates with both ADAM22 and ADAM23 to be properly processed with Endo H-resistant N-glycans in HEK cells. Moreover, this mutation does not allow the molecule to be anchored at the AIS suggesting a loss of interaction with other unknown partners specifically expressed in that site. LGI1 is known to interact with NgR1 that may enhance the association of LGI1 with ADAM22 (Thomas et al., 2016). Whether any mutation in LGI1 linked with epilepsy may induce defect in NgR1 binding is unknown (Thomas et al., 2016). ADAM11 is also known as a ligand for LGI1 (Sagane et al., 2008) that plays a critical role for localizing the Kv1 channels at the presynaptic terminals of cerebellar basket cells (Kole et al., 2015). There is no indication that these ligands of LGI1 may be localized at the AIS. Thus, the pathogenic mechanisms for the LGI1^S473L^ mutation may rely both on its defective binding to one of its receptors and to its mistargeting into axonal subcompartments.

LGI1 has been proposed to form a trans-synaptic bridge through its binding with ADAM22 and ADAM23 expressed at the post- and pre-synapse, respectively. ADAM22 and ADAM23 are partitioned into synaptic fractions depending on LGI1 (Fukata et al., 2010). LGI1/ADAM22 binds PSD-95 and consequently may stabilize the AMPAR and stargazin complex, regulating synaptic strength at the excitatory synapse (Fukata et al., 2006). Interestingly, LGI1 seems to be required for the synaptic localization of ADAM22 and ADAM23 as indicated by immunohistochemical and biochemical analyses of LGI1-deficient mice (Fukata et al., 2010). Conversely, in *adam22^-/-^* and *adam22^-/-^* mice, the neuropil staining of LGI1 is lower in most hippocampal regions indicating that these molecules are interdependent for their distribution at the synaptic neuropil (Yokoi et al., 2015). Our results show that LGI1 may associate with ADAM22 or ADAM23 early along the secretory pathway at the ER level to be sorted with either ADAM molecule in axonal transport vesicles. We observed that vesicles containing ADAM22 and ADAM23 either alone or co-localized with LGI1 are axonally transported through the AIS as reported for axonal cargoes (Al-Bassam et al., 2012; Petersen et al., 2014). The vesicular axonal transport of ADAM23 was predominantly oriented in the anterograde direction whereas ADAM22 was preferentially transported in the retrograde direction. Such a differential transport may promote a distinct pre- and post-synaptic distribution at the steady state. Therefore, the interdependance of LGI1 and ADAM proteins for their synaptic distribution may be based on their association during axonal transport. As we observed for LGI1 and ADAM proteins, the axonal transport of Kv1.2 subunits associated with the accessory Kvß2 subunits has been reported to occur both in the anterograde and retrograde directions with similar velocities (Gu and Gu, 2010). The axonal vesicular transport of Kv1.2 is facilitated by Kvß2 and depends on kinesin KIF3A and KIF5B (Gu and Gu, 2010; Rivera et al., 2007). Whether LGI1 and ADAM proteins may traffic together with Kv1 channels as a preformed complex and using identical molecular motors deserves further investigations.

The composition of the Kv1 complex and the mechanisms regulating its recruitment at the AIS are still elusive. The analysis of the Caspr2 interactome in hippocampus indicates its association with TAG-1 and Kv1, but also with ADAM22 and LGI1 (Chen et al., 2015). Using co-immunoprecipitation experiments from transfected HEK cells, we determined that Caspr2 interacts through its ectodomain with ADAM22 and ADAM23, whereas TAG-1 only precipitated with ADAM22. Moreover, the ADAM proteins can be sorted together with Caspr2 or TAG-1 in axonal transport vesicles. We recently showed that TAG-1 is enriched at the AIS whereas Caspr2 is uniformly expressed along the axon of cultures hippocampal neurons (Pinatel et al., 2017). Similarly, here we noticed that ADAM23 is enriched at the AIS when co-transfected with TAG-1, but not when co-transfected with Caspr2. Therefore the ADAM family may play a pivotal role for the interdependent distribution of the different sets of CAMs associated with Kv1 at the AIS. Our results indicate that the focus on AIS might be relevant for the further dissection of the pathogenic mechanisms implicating LGI1 in epilepsy.

## Materials and Methods

### Constructs

The pCDNA3-Caspr2-HA construct encodes human Caspr2 with the HA epitope inserted downstream the signal peptide between the residues Trp26 and Thr27 (Bel et al., 2009). The Caspr2-HA deleted construct Caspr2Δcyt (stop codon at aa 1285) was described previously (Pinatel et al., 2015). NrCAM-GFP was previously described (Falk et al., 2004). The human TAG-1-GFP and Caspr2-GFP constructs with GFP downstream of the signal peptide were described (Pinatel et al., 2015). Caspr2-mCherry was generated by insertion into the EcoRI-BamHI sites of pmCherry-N1. Plasmids encoding human LGI1, ADAM22, ADAM23 were purchased from Origene. LGI1-GFP was generated by insertion in pEGFP-N3 (Pinatel et al., 2015). The R407C, S473L and R474Q missense mutations of LGI1-GFP were generated using QuickChange II mutagenesis kit (Stratagene). ADAM22-mCherry and ADAM23-mCherry were generated by insertion into the NheI-KpnI sites of pmCherry-N1. PCR amplified products were verified by sequencing (Genewiz).

### Western blot and immunoprecipitation

HEK cells were co-transfected with Caspr2-HA, Caspr2Δcyt-HA, LGI1-GFP or TAG-1-GFP, and ADAM22-mCherry or ADAM23-mCherry. Cells were lyzed for 30 min on ice with 50 mM Tris, pH 7.5, 1% NP-40, 10 mM MgCl2 and protease inhibitors, centrifuged at 4°C for 15 min at 15,000 rpm. After preclearing for 1 h at 4°C with Protein G-Sepharose, supernatants were immunoprecipitated overnight at 4°C with protein G-agarose coated with rabbit anti-mCherry (RFP) antibody (2 μl) and mouse anti-GFP IgG (1 μg), or rat anti-HA IgG (1 μg). The beads were washed twice with 50 mM Tris pH 7.4, 150 mM NaCl and 1% NP-40, twice in 50 mM Tris, 150 mM NaCl and twice in 50 mM Tris. Immune precipitates were analyzed by immunoblotting with anti-HA, anti-GFP, or anti-mCherry antibodies. Blots were developed using the ECL chemiluminescent detection system (Roche). HEK cells were transfected with LGI1-GFP or LGI1^S473L^-GFP alone or co-transfected with ADAM22-mCherry or ADAM23-mCherry. The LGI1 proteins were immunoprecipitated from the lysates using mouse anti-GFP, eluted from Protein G-Sepharose in Tris 50 mM pH6, 0.2% SDS, 1%β-mercaptoethanol, 0.5% Triton-X100, 1 mM EDTA and protease inhibitors and incubated for 3 h with Endoglycanase H (3 mU/μl) or PNGase F (1 U/μl) (Roche).

### Antibodies and immunofluorescence staining

The chicken anti-MAP2 (ab5392) and the goat anti-GFP (ab 5450) antibody were purchased from Abcam, the rat anti-HA mAb (clone 3F10) and the mouse anti-GFP (cat. 11 814 460 001) from Roche, the rabbit anti-RFP (anti-mCherry, code 600-401-379) antiserum from Rockland, the rabbit anti-ADAM23 (cat PA5-30939) from ThermoScientific. The mouse anti-AnkyrinG (clone N106/36), anti-ADAM22 (clone N57/2) anti-LGI1 (N283/7), anti-pan neurofascin (clone A12/18) and anti-Kv1.2 (clone K14/16) mAbs were obtained from the UC Davis/NIH NeuroMab facility. The mouse anti-TAG-1 1C12 was a gift from Dr. D. Karagogeos. AlexaFluor488-, 568- and 647-conjugated secondary antibodies were obtained from Molecular Probes. Immunostaining for Caspr2-HA, LGI1-GFP, TAG-1-GFP, and ADAM23 were performed on live cells with antibodies diluted 1:500 in culture medium for 30-60 min. Cells were fixed with 4% paraformaldehyde in PBS for 10 min and permeabilized with 0.1% Triton-X100 for 10 min. Immunofluorescence staining was performed using chicken anti-MAP2 (1:10,000) and mouse anti-AnkyrinG (1:100) antibodies, and with secondary antibodies diluted in PBS containing 3% bovine serum albumin. After washing in PBS, cells were mounted in Mowiol (Calbiochem). Since the direct fluorescence of LGI1-GFP was very low after fixation with paraformaldehyde, the detection of LGI1-GFP at the neuronal surface was performed using live immunolabelling with anti-GFP antibody and either Alexa568 or Alexa488 secondary antibody.

### Cell culture

Cell culture media and reagents were from Invitrogen. HEK-293 cells recently authenticated and tested for contamination were grown in DMEM containing 10% fetal calf serum and were transiently transfected using jet PEI (Polyplus transfection, Ozyme). Primary hippocampal cell cultures were prepared from embryonic day 18-Wistar rats. Hippocampi were collected in Hanks’ balanced salt solution, dissociated with trypsin and plated at a density of 1.2 10^5^ cells/cm^2^ on poly-L-lysine coated coverslips. The hippocampal neurons were cultured in Neurobasal supplemented with 2% B-27, 1% penicillin-streptomycin and 0.3% glutamine in a humidified atmosphere containing 5% CO2 at 37°C. Hippocampal neurons were transfected using Lipofectamine 2000 with Caspr2-HA Caspr2-GFP, TAG-1-GFP, LGI1-GFP, NrCAM-GFP, ADAM22-mCherry and ADAM23-mCherry or co-transfected with two of these constructs at DIV8 or DIV13. All animal experiments were carried out according to the European and Institutional guidelines for the care and use of laboratory animals and approved by the local authority (laboratory’s agreement number D13-055-8, Préfecture des Bouches du Rhône).

### Confocal microscopy and image analysis

Image acquisition was performed on a Zeiss laser-scanning microscope LSM780 equipped with 63 X 1.32 NA oil-immersion objective. Images of GFP or mCherry or AlexaFluor-stained cells were obtained using the 488 nm band of an Argon laser and the 568 nm and 647 nm bands of a solid state laser for excitation. Fluorescence images were collected automatically with an average of two-frame scans. Quantitative image analysis was performed using Image-J on confocal sections (10-20 neurons in each condition). The fluorescence intensity was measured in 2 regions of interest (AnkyrinG-positive AIS and axon) using identical confocal parameters. Regions corresponding to AIS were manually selected on AnkyrinG images and reported on other channels for intensity measurements. All intensities were corrected for background labelling using the Zen software (Zeiss). Statistical analysis was performed using Statview or GraphPad Prism software. The data normal distribution was tested using d’Agostino and Pearson’s test. For multiple group comparisons, we used one-way ANOVA followed by Fisher’s test. Instead, the non-parametric Mann-Whitney test was used when the assumption of normality was not possible.

### Imaging vesicle transport

Coverslips with neurons were loaded into a sealed heated chamber in imaging medium (Hank’s balanced salt solution pH7.2 with 10 mM HEPES and 0.6 % glucose). Recordings were made 18 h after transfection. The axons were selected on the basis of their much greater length by comparison with dendrites. Live immunostaining using AlexaFluor647-conjugated Neurofascin186 was performed to visualized the AIS. Vesicle transport was imaged using Zeiss laser-scanning microscope equipped with 63 X 1.32 NA oil-immersion objective and 37°C heating chamber. Dual-color recordings were acquired using simultaneous excitation with 488 (2-4 %) and 561 lasers (1-2 %), and GaSP PMT1 for 499-551 and PMT2 for 569-735 detections (562x240 pixels, average 2, open pinhole, 1.5 s scanning time, streamed time-lapse recording during 3-9 min). Kymographs were generated using ImageJ software and contrast inverted so that the fluorescent vesicles corresponded to dark lines. Overlapping transport events were analyzed and the velocity measured.

## Acknowledgements

We wish to thank Marie-Pierre Blanchard of the CRN2M imaging core facility for help with time-lapse recording and image analysis. We are grateful to Michael Seagar, Jérôme Honnorat, Laurence Goutebroze and Christophe Leterrier for helpful discussions. We thank the UC Davis/NIH NeuroMab facility. This work was supported by the Association pour la Recherche sur la Sclérose en Plaques (ARSEP) to CFS.

